# The signal sequence of yeast killer toxin K2 confers producer self-protection and allows conversion into a modular toxin-antitoxin system

**DOI:** 10.1101/2023.10.24.563802

**Authors:** Rianne C. Prins, Sonja Billerbeck

## Abstract

Some antimicrobial proteins secreted by yeast, known as yeast killer toxins, also target the producer species itself, necessitating a means of self-protection. Intriguingly, the M2 dsRNA killer virus in *Saccharomyces cerevisiae* contains a single open reading frame (ORF) that encodes both the pore-forming killer toxin K2 as well as a cognate immunity factor. Here, a systematic deletion screen reveals that expression of a 49-amino acid N-terminal peptide from this ORF is both necessary and sufficient for immunity and that the K2 toxin and this 49-residue immunity peptide can be functionally split into a modular toxin-antitoxin system. Further, the immunity peptide exhibits characteristics of a signal peptide and we thus propose that the K2 signal peptide serves a dual function: 1) Toxin targeting into the secretory pathway, and 2) establishing self-protective immunity. This case further implies that (signal) peptides form a potential source for antimicrobial resistance.

## Introduction

Yeast killer toxins (YKTs) are proteinaceous toxins secreted by killer yeast. They are widespread and exhibit diverse mechanisms of killing and target cell spectra^1–4^. The secretion of YKTs can provide a growth advantage in a given environmental niche by eliminating potential invaders or competitors^5^. However, several YKTs do not solely target other microbial species, but also affect the producer strain itself. Consequently, the toxin-producer requires mechanisms of self-protection, also referred to as immunity. This phenomenon bears resemblance to self-selecting bacterial toxin-antitoxin systems, where offspring that has lost the toxin-antitoxin system is eliminated^6^.

Killer toxins and their cognate immunity factors are genetically encoded in various ways. Some YKTs are encoded on plasmids and accompanied by dedicated immunity genes, exemplified by zymocin from *Kluyveromyces lactis*, PaT from *Pichia acacia,* or DrT from *Debaryomyces robertsiae*^7,8^. Interestingly, other YKT-immunity pairs can arise from single open reading frames (ORF). This is observed in several strains of *Saccharomyces cerevisiae*, more specifically those that harbor the cytoplasmic double-stranded RNA (dsRNA) viruses M1, M2, M28 or Mlus^4,9^. These M-viruses are small satellite viruses (1.6-2.3 kb) that are dependent on a larger L-A dsRNA helper virus (4.6 kb) for encapsidation and replication and are vertically transmitted^4,9,10^. Their compact genomes contain a sole ORF that encodes a killer toxin (K1, K2, K28, Klus) and a corresponding immunity factor.

While the immunity mechanism for K28 has been elucidated and requires the intracellular K28 precursor that deactivates internalized K28 toxin by complex formation and subsequent ubiquitination and proteolytic degradation^11^, those used by the other killer viruses are less evident. In this study, we investigated the immunity factor of the K2 toxin. It is known that the K2 killer toxin initially interacts with β-1,6-glucans in the target cell wall^12^, and subsequently translocates to the plasma membrane where it likely interacts with a plasma membrane receptor^13^, and disrupts plasma membrane integrity^12,14,15^ by inducing ion leakage similar to the ionophore toxin K1^16^. A K2-specific immunity factor prevents this toxicity in the producer cell. Previous mutagenesis of the K2 precursor indicated that several regions from the N- to the C-terminus are important for the K2 immunity factor^17,18^, but its properties remain unclear.

Here, we show with a plasmid-encoded K2 system that expression of an N-terminal 49-residue peptide of the K2 precursor provides wild-type level protection to the K2 toxin. Our data suggest that this peptide also encodes the N-terminal signal peptide of K2 and as such has dual functionality – a rather rare phenomenon for signal peptides: it acts as the immunity factor after targeting the K2 toxin to the secretory pathway. When replacing this dual-function N-terminal signal peptide with a canonical secretion signal peptide, we show that K2 can be functionally split into two ORFs, one encoding the immunity and one the toxin. This resembles a prokaryotic system of a bacteriocin gene accompanied by a separate gene encoding a cognate immunity protein, but those have not been reported to exist in *S. cerevisiae*.

## Results

### 1. A systematic deletion scan localizes the immunity factor to the K2 N-terminus

To gain insights into how the immunity factor is encoded within the 363-codon K2 ORF, we created a series of systematic sequential deletions within the K2 gene (**Figure 1A**). We hypothesized that constructs carrying deletions within regions unrelated to immunity would still support cell growth in the presence of externally added K2 toxin, whereas constructs with deletions that impacted immunity would not provide protection and thus cells would not be able to grow in the presence of externally added K2 toxin. Each deletion involved the substitution of 14-27 codons with a SbfI restriction site (CCTGCAGGG) – which simultaneously generated a collection of backbones suitable for future cloning purposes. The SbfI-insertion added three in-frame amino acids, namely proline, alanine, and glycine (P-A-G). Given the potential for these constructs to exhibit suicidal phenotypes, all K2 variants were cloned downstream of a galactose-inducible promoter on a high-copy pRS423-type plasmid. K2 expression could thus be repressed in the presence of glucose and induced in the presence of galactose.

**Figure 1.**
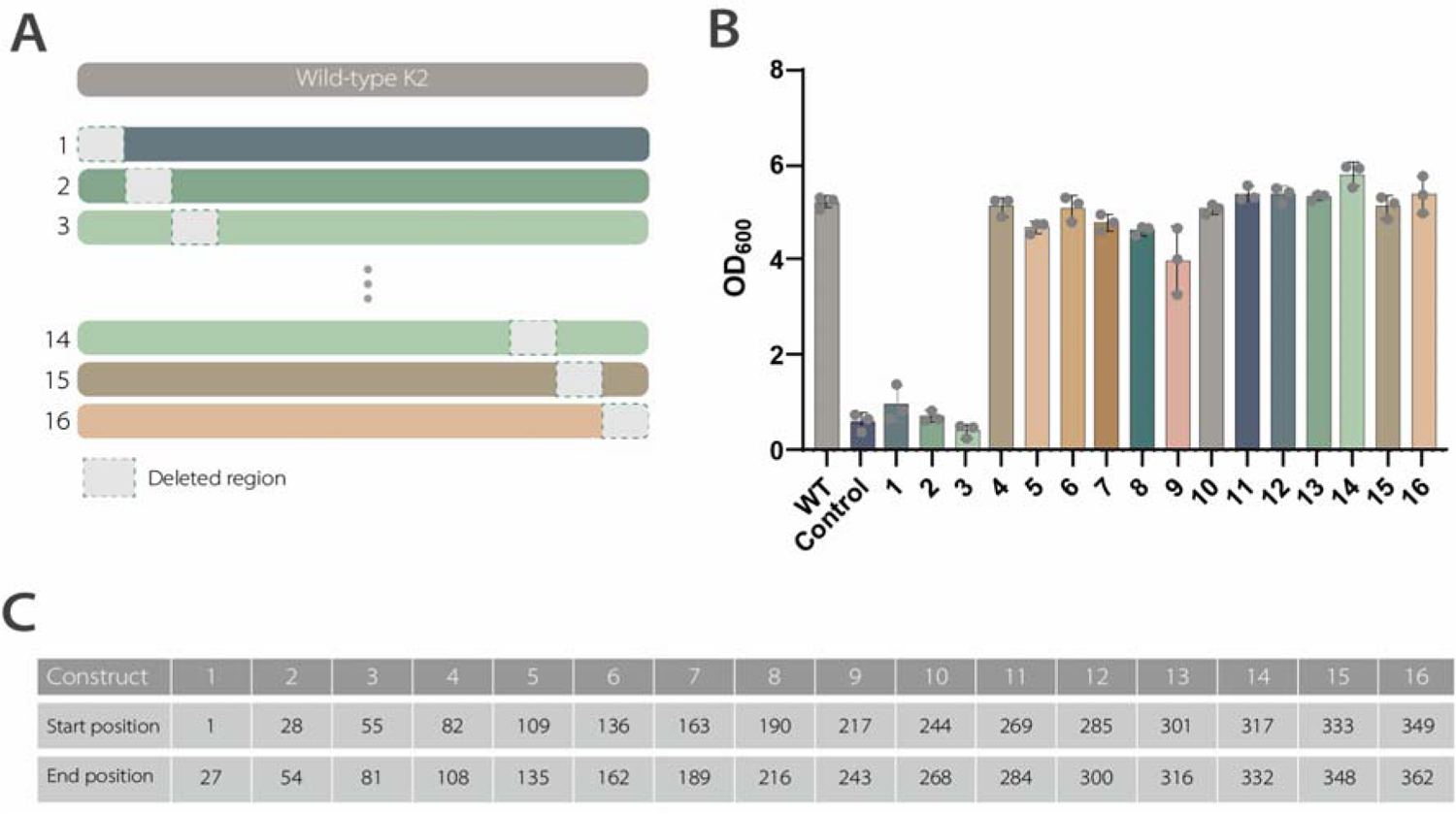
Significance of the N-terminal region in K2 immunity. **A)** Overview of the 16 deletion constructs, each lacking a region (14-27 codons) within the K2 ORF (not drawn to scale). The excised segments were replaced with an in-frame SbfI restriction site encoding Pro-Ala-Gly and the genes placed behind a GAL1 galactose-inducible promoter on a pRS423-type vector. Following yeast transformation, three transformants were cultivated overnight in inducing media containing 1% galactose, before being transferred to fresh inducing media supplemented with 10 a.u. of K2 toxin. **B)** The OD_600_ in toxin presence was recorded in a plate reader over the course of 24 hours. The bars represent the mean of the final OD_600_ values of biological triplicates, with error bars indicating ± 1 standard deviation. Individual replicate values are displayed as dots. WT: Wild-type K2, Control: Empty vector. **C)** The table provides information regarding the first and last deleted codons of the modified regions of the systematic deletion constructs. Note that the first construct also omits the start codon at position 1.

We assessed the immunity function using a standardized workflow: *S. cerevisiae* was transformed with one of the respective 16 constructs, and three resulting colonies were arrayed into 96-well microtiter plates filled with non-inducing liquid media. The cultures were allowed to reach comparable levels of overnight growth saturation. Following this, the cultures were diluted at a 1:20 ratio into protein-expression media supplemented with 1% galactose and incubated for 24 hours. Throughout this incubation period, growth was monitored to determine if the mere expression of the constructs had any impact on cell growth, and this timeframe in addition allowed potential cellular adaptation to develop immunity prior to exposure to high levels of externally added toxin. In the last step, cultures were diluted into assay plates with media containing 1% galactose and 10 a.u. of externally added K2 toxin. Growth was once again recorded by OD_600_ readings over a 24-hour span. The use of 1% galactose ensured high K2 expression and served to define initial on/off immunity phenotypes.

The externally added toxin was prepared by concentrating the supernatant of a K2 toxin producer strain (see Methods) and 10 a.u. K2 toxin was established as a concentration that reliably killed non-immune cells while enabling normal growth of immune cells in our assay system (**Figure S1A**).

All deletion constructs led to similar growth in the absence of externally applied K2 toxin (**Figure S1B**). When challenged with 10 a.u. of K2 toxin, 13 out of the 16 constructs were able to sustain growth levels similar to the wild-type, despite featuring deletions across a substantial portion of the ORF (**Figure 1B**). This observation strongly implies that the immunity factor remains highly preserved in these 13 constructs. In contrast to a study in 2002^18^ that proposed the involvement of the α- and β-subunits in immunity, in addition to the N-terminal region, we observed no large effect on K2 immunity when deletions were introduced in the respective regions (constructs 7, 8, 15 and 16). Remarkably, the three constructs that led to a loss-of-resistance phenotype (1, 2, and 3) contained deletions spanning amino acid positions 1 to 81 (**Figure 1C**). This finding indicates that the key information regarding the K2 immunity factor is strongly associated with this N-terminal region.

### 2. The K2 N-terminus also encodes a potential signal peptide for secretion

Given that the N-termini of secreted proteins typically harbor signal sequences responsible for directing the downstream protein into the secretory pathway, we investigated the possibility that the K2 signal peptide in addition serves as the immunity factor. We used specialized prediction tools to predict the signal sequence of the K2 toxin (i.e. SignalP-6.0^19^, TargetP-2.0^20^) but interestingly they failed to identify a signal peptide in the K2 precursor **(Figure S2A)**. Given K2 is a secreted protein and thus requires a signal sequence for secretion, we manually searched the N-terminus for known features of a signal peptide. Signal sequences in *S. cerevisiae* are typically 18-24 residues long, and they feature a hydrophobic alpha-helical region (H-region), flanked by a positively charged N-terminal region (N-region) and a C-terminal region (C-region) that potentially contains a recognition sequence for the signal peptidase^21^ (Figure 2B).

**Figure 2.**
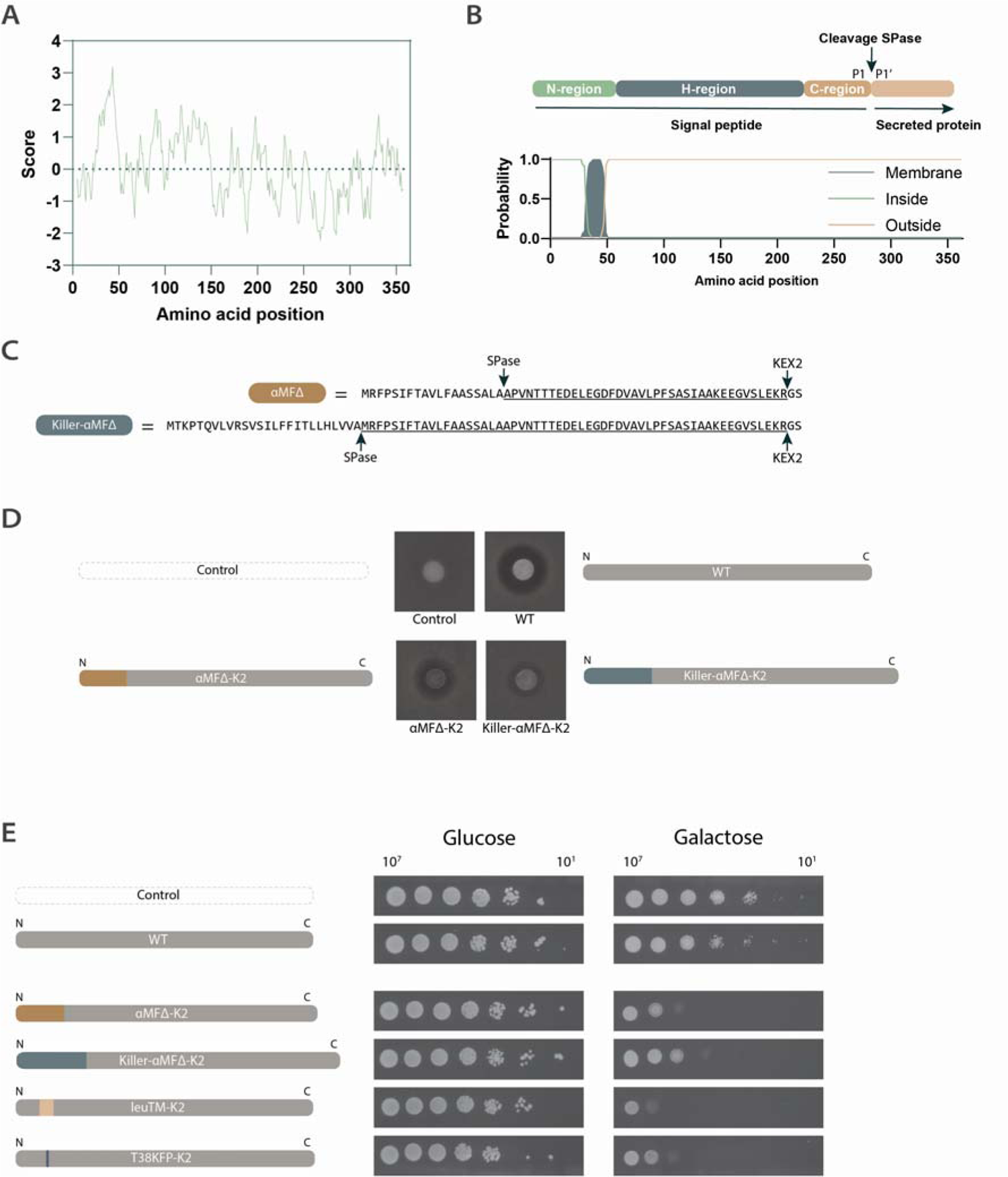
The crucial role of the predicted K2 signal peptide in immunity. **A)** Kyte and Doolittle hydropathy plot^22^ of the K2 precursor. **B)** Scheme of the signal sequence domains (top): An N-region, typically containing positively charged residues, a hydrophobic H-region, and a C-region housing the signal peptidase cleavage site. The bottom panel depicts the predicted H-region in the K2 precursor. **C)** The two signal peptides αMFΔ and killer-αMFΔ contain a pre-region subject to signal peptidase cleavage, and a pro-region (underscored) processed by KEX2 proteases after the dibasic recognition site. The used cloning approach^29^ results in the addition of linker residues Gly-Ser to the downstream protein sequence. **D)** Toxicity assay based on the formation of a zone of inhibition of producer spots on a lawn of sensitive cells. A strain with a *pbs2* gene deletion was employed as a sensitive strain due to its high sensitivity to K2 toxin^30^, thereby enhancing the visual distinction of halo sizes. The αMFΔ and killer-αMFΔ signal peptides were used to secrete K2 without its native signal peptide. Plates were incubated for 3 days at 25°C before imaging. **E)** *S. cerevisiae* cells were transformed with the respective variant constructs. Transformant cultures were spotted in 10-fold dilutions (10^7^ to 10^1^ cells) on non-inducing (glucose, left) or inducing plates (1% galactose, right). Cell growth was imaged after 3 days (glucose: 30°C, galactose: 25°C) and representative results are shown. The K2 wild-type sequence is depicted in grey. Sequence alterations are indicated by colored blocks. SPase: signal peptidase. WT: Wild-type K2 producer, Control: Empty vector, αMFΔ and Killer-αMFΔ: Signal peptides based on the α-mating factor, LeuTM-K2: Poly-leucine transmembrane region from 30 to 47, T38KFP-K2: Mutation of T at position 38 to KFP.

First, we searched for a hydrophobic domain: A hydropathy plot^22^ of the K2 precursor indicated a distinctive hydrophobic domain within the first N-terminal 50 residues of K2 (Figure 2A). This was further substantiated by transmembrane domain and signal peptide prediction tools such as Phobius, TMHMM-2.0, and deepTMHMM^23–25^, with a consensus hydrophobic domain between residues 31 and 47 (Figure 2B). In the downstream sequence, multiple putative signal peptidase cleavage sites are present (see **Supplementary Note 1**). On the N-terminal side of this H-region, K2 encodes two positively charged residues (arginine), aligning with characteristic features of N-regions. We hypothesized that failure to predict the signal peptide with bioinformatic tools was likely the result of an unusually lengthy N-region of 30 residues rather than the canonical ∼5 residues^26^. Notably, SignalP-6.0 does recognize a signal sequence at the N-terminus of the K2 precursor with high confidence when the N-region is shortened by the removal of 17 N-terminal residues *in silico* (**Figure S2A and B**). An AlphaFold^27^ structure prediction of the K2 precursor is further consistent with the N-terminal signal peptide domain predictions **(Figure S2C-E)**: The H domain is helical and the signal peptide displays a separate N-terminal domain, consistent with the expected characteristics of signal peptides that are not part of the mature secreted protein.

### 3. Replacing the dual-function N-terminal signal peptide with a canonical signal peptide allows to functionally split the toxin from its immunity factor

Given our results indicated that the immunity factor was potentially located within the N-terminal signal peptide we wanted to determine the functional modularity of the K2 toxin and the immunity factor of K2. We therefore conducted a substitution experiment, replacing the original predicted signal sequence region (here residues 1 to 54, see **Supplementary Note 1**) with an unrelated signal sequence to test whether this would result in a functional toxin while eliminating immunity, thereby inducing a suicidal phenotype. For this purpose, we selected two signal peptides from the Pichia cloning toolkit^28,29^ and fused these to residues 54-362 of the K2 precursor. The chosen signal peptides are derived from the *S. cerevisiae* α-mating factor signal peptide (αMF), a widely used secretion signal in yeast. A shortened version of this signal peptide (αMFΔ), comprises a pre-region that is cleaved by signal peptidases, followed by a pro-region that undergoes processing at a KEX2 protease site within the secretory pathway (Figure 2C). The αMF lacks a strong transmembrane helix prediction and facilitates translocation *via* the posttranslational translocation pathway. Combinations of this sequence with additional N-terminal pre-regions have demonstrated enhanced efficiency, and we included an αMFΔ peptide with an N-terminal pre-region originating from the K1 killer toxin, an element available in the toolkit (killer-αMFΔ).

Functional toxin expression was verified by the formation of a zone of inhibition on a background lawn of a sensitive strain. Indeed, K2 toxin expression directed by αMFΔ and killer-αMFΔ resulted in halo formation indicating secretion of active toxin (Figure 2D). However, it was also apparent that the producer strains exhibited less dense spots, implying a reduced cell population. This phenotype could be due to self-killing, attributed to the lack of the original N-terminal signal peptide required for self-protection.

To observe the potential suicidal phenotype in more detail, we conducted a spot assay of culture dilutions (Figure 2E). Strains harboring the αMFΔ-K2 and killer-αMFΔ-K2 constructs show a normal growth phenotype under non-inducing conditions (glucose), but clearly become suicidal in inducing conditions (galactose). αMFΔ-K2, the construct with the larger halo, also shows a larger degree of self-killing compared to killer-αMFΔ-K2.

We also included two additional constructs in this assay. One of them features the previously reported substitution of position T38 by K-F-P (T38KFP-K2) known to result in a suicidal phenotype^17^. The other construct carries a replacement of the predicted transmembrane domain by a highly hydrophobic poly-leucine repeat (spanning residues 30 to 47) (LeuTM-K2). Both of these constructs demonstrate strong suicidal phenotypes, with the poly-leucine variant showcasing the highest degree of self-killing across all tested constructs (Figure 2E). Collectively, these observations indicate that the killer toxin can be secreted independently from its immunity factor and that specific sequence-function information for immunity exists within the predicted K2 signal peptide H-region.

### 4. A 47-aa N-terminal peptide is necessary and sufficient for toxin immunity

Following the findings that the toxin could be functionally expressed when fused to a different signal sequence, leading to suicidal phenotypes, and the preceding data emphasizing the critical role of the N-terminus in conferring immunity, a key question remained: Can expression of an N-terminal peptide alone also suffice to provide immunity?

To address this question, we engineered a series of C-terminally truncated variants, ranging from 108 amino acids down to only 15 N-terminal amino acids (Figure 3A). Transformants expressing these constructs showed no growth phenotype in inducing media alone (**Figure S3**), but when challenged with 10 a.u. of K2 toxin, those containing at least 47 N-terminal amino acids demonstrated immunity, whereas those with 43 amino acids or less exhibited a loss-of-resistance phenotype (Figure 3B). Thus, an N-terminal peptide of 47 amino acids is not only indispensable, moreover, it has the capacity to independently confer resistance to the K2 toxin.

**Figure 3.**
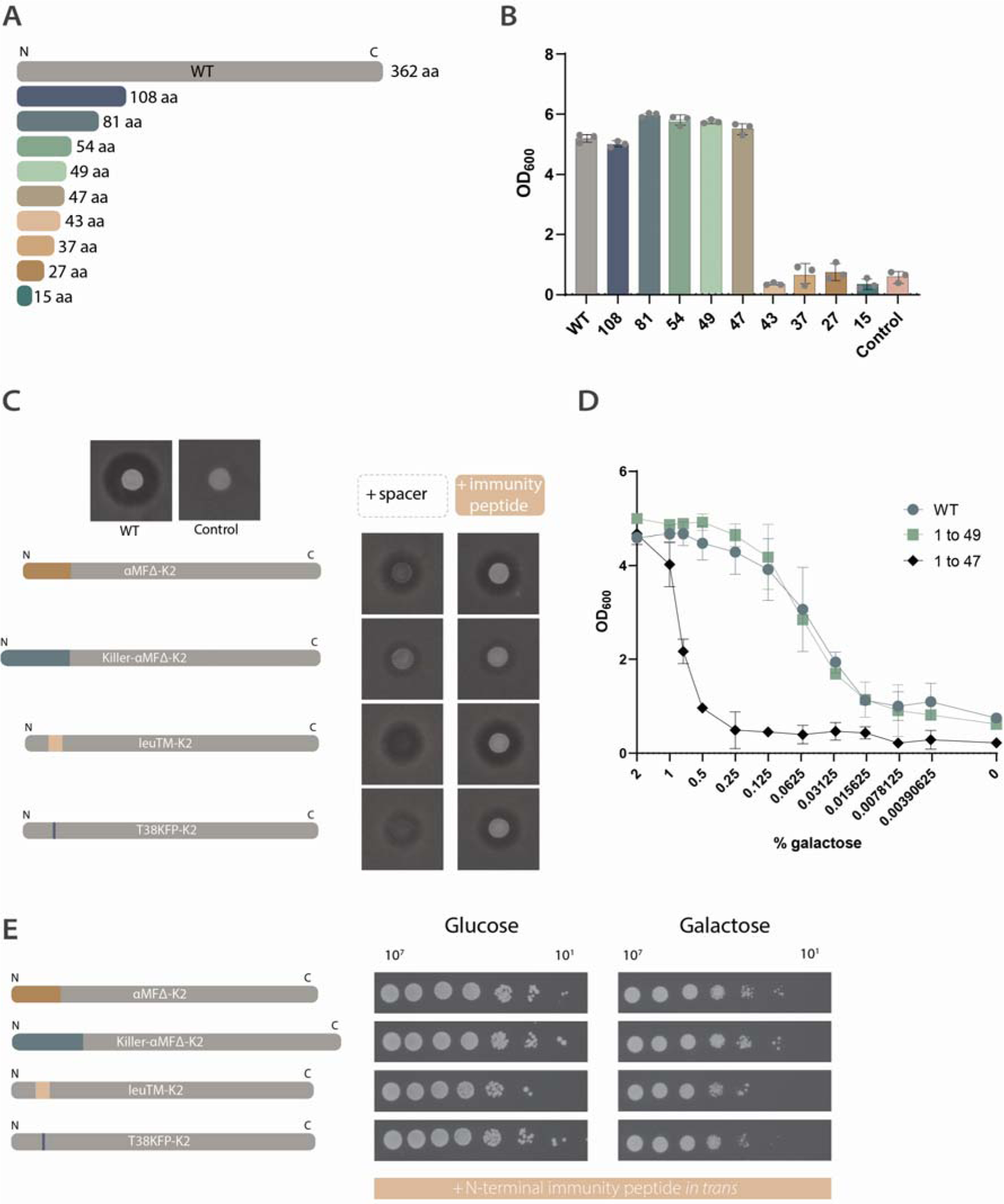
A 47-aa N-terminal peptide suffices to confer K2 resistance and allows the building of a synthetic toxin-antitoxin system. **A)** Overview of the tested N-terminal protein sizes, from the wild-type (362 aa) down to 15 aa. **B)** The OD_600_ in the presence of 1% galactose and 10 a.u. of K2 toxin was recorded in a plate reader over 24 hours. The final OD_600_ values are depicted. Bars represent the mean of biological triplicates, with error bars indicating ± 1 standard deviation, and individual replicate values are displayed as dots. WT: Wild-type K2, Control: Empty vector. **C)** Halo assays of the suicidal constructs complemented with an empty ‘spacer’ plasmid or a plasmid carrying the 47-aa immunity peptide. Note that a few of these images are duplicated from Figure 2 to facilitate comparison of the spots. **D)** Cells producing either wild-type K2, or peptides consisting of residues 1-49 or 1-47 were cultivated in the presence of varying concentrations of the inducer galactose, and exposed to 10 a.u. of K2 toxin. The final OD_600_ values after 24 hours of incubation are displayed, as the mean of biological triplicates, with error bars indicating ± 1 standard deviation. **E)** Transformants were spotted in 10-fold dilutions (10^7^ to 10^1^ cells), either on non-inducing (glucose, left) or inducing (1% galactose, right) plates. Cell growth was imaged after 3 days of incubation (glucose: 30°C, galactose: 25°C) and representative results are shown (*n*=2). K2 wild-type sequence is colored grey. Changes in sequence are indicated by colored blocks. WT: wild-type K2 producer, Control: Empty vector, αMFΔ and Killer-αMFΔ: signal peptides based on the α-mating factor, LeuTM-K2: poly-leucine transmembrane region from 30 to 47, T38KFP-K2: mutation of T at position 38 to KFP.

When we investigated immunity across different protein expression levels, the concentration-dependent nature of the immunity factor showed (Figure 3D). Upon induction with varying levels of galactose, while maintaining a constant toxin challenge of 10 a.u., we observe distinct immunity profiles between the peptides comprising residues 1-47 and 1-49. Although both peptides confer similar, robust immunity when highly expressed, peptide 1-47 exhibits weaker immunity with reduced expression levels compared to 1-49. Notably, cells expressing the peptide 1-49 mirror the performance of those expressing wild-type K2. This observation suggests that residues K48 and S49, while less vital at higher expression levels, become crucial for effective protection at lower expression levels.

It has been determined that approximately 450 molecules of K2 toxin can bind to a single cell^12^. Given the concentration-dependent nature of the immunity protein, this implies that a certain number of immunity proteins are necessary to achieve complete resistance to the K2 toxin. These immunity proteins may play a role in interacting with the toxin molecules or blocking a receptor that only becomes saturated with a certain number of immunity proteins.

### 5. The immunity peptide and the K2 toxin can be functionally co-expressed as a synthetic toxin-antitoxin system

The fact that both the toxin and the immunity factor can be expressed separately, implies the potential of this system for constructing a modular toxin-antitoxin system in *S. cerevisiae*. As such, we cloned the 47-aa peptide under control of a galactose-inducible promoter on a high-copy pRS426-type plasmid and co-transformed it together with the various K2 variants with exchanged signal peptides. We then tested for a growth phenotype on inducing media (galactose) using the same serial dilution spot assay as described above. In fact, the *in trans* expression of the small immunity peptide within the previously suicidal strains demonstrated rescue of the suicidal phenotype and restored growth comparable to cells carrying wild-type K2 (Figure 3C and D).

### 6. Further characterization reveals a frame within the immunity peptide critical for function

The constructs tested so far indicate that residues within the predicted transmembrane region are important, as well as a certain number of C-terminal residues. To further determine the requirements for immunity, we also created a series of 8 N-terminally truncated constructs (Figure 4A). We systematically deleted residues starting at the N-terminus, up to residue 24, while keeping an initiator methionine at position 1.

**Figure 4.**
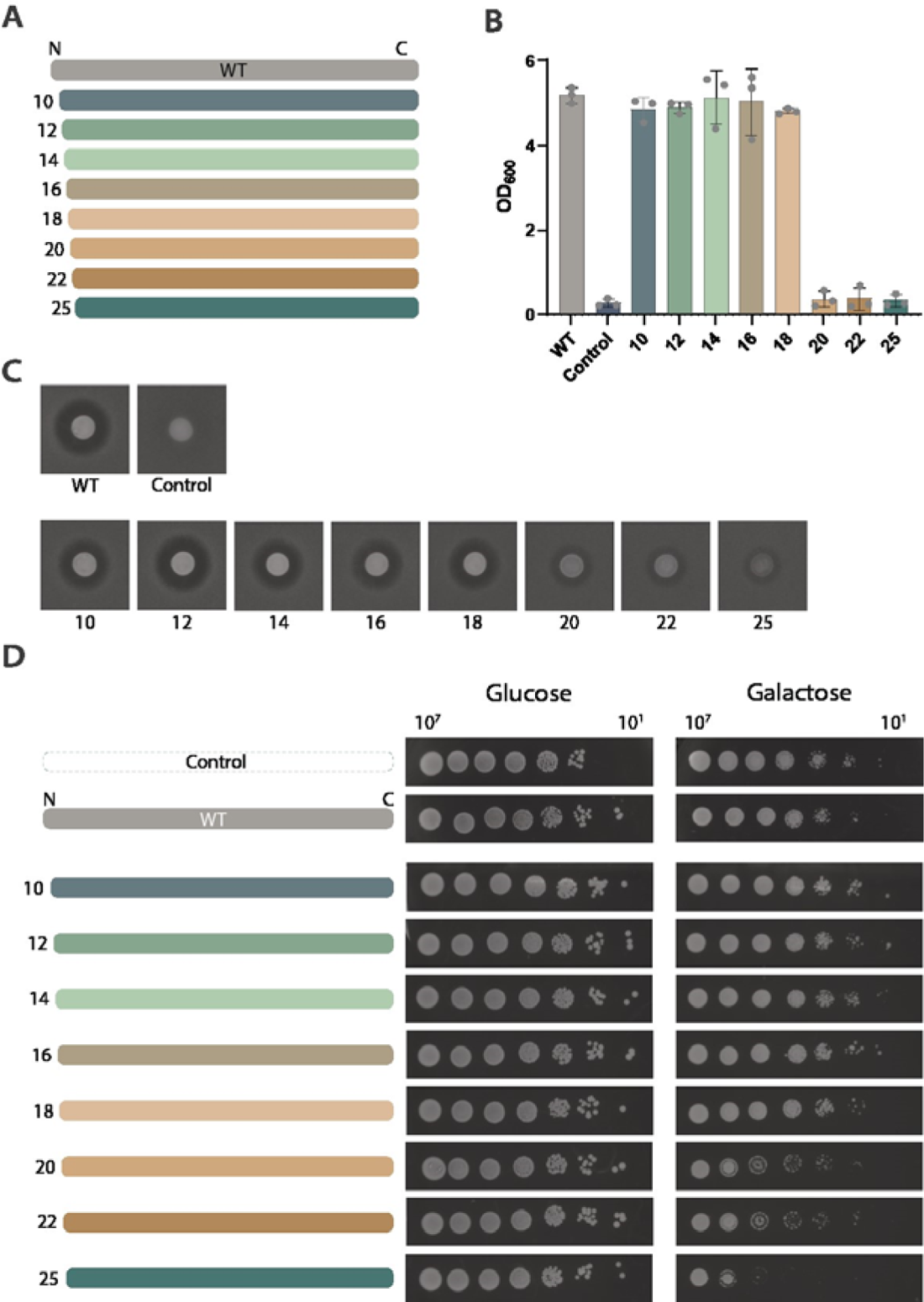
Systematic N-terminal truncations point to critical residues within the immunity peptide. **A)** Visualization of the tested constructs. The number indicates the amino acid position in the wild-type sequence that is first expressed, after the added initiator methionine. **B)** Yeast transformants were assayed in the presence of 10 a.u. K2 toxin (and 1% galactose). Final OD_600_ values after 24 hours were recorded. Bars represent the mean of three biological replicates with error bars representing +-1 standard deviation. Individual replicates are depicted as dots. **C)** To assay active toxin production, the respective strains were spotted (10^7^ cells) on top of agar plates containing a sensitive background strain and incubated for 3 days at 25°C before imaging zones of inhibition. **D)** To assay potential suicidal phenotypes, strains were spotted in logarithmic dilutions (10^7^ to 10^1^ cells) onto agar plates with non-inducing (glucose, left) or inducing (1% galactose, right) conditions. Cell growth was imaged after 3 days of incubation (glucose: 30°C, galactose: 25°C) and representative results are shown (*n*=2). WT: Wild-type K2, Control: Empty vector.

The resulting constructs were assayed for immunity in liquid media supplemented with 10 a.u. of toxin as described above. Construct expression did not affect growth (**Figure S4A)**, but when the cells were challenged with 10 a.u. of toxin, only the constructs that at least contained the sequence from residue 18 onwards conferred immunity (Figure 4B). Sequences starting at position 20 lacked sufficient protection. Interestingly, the constructs that lead to a loss-of-resistance phenotype still support functional K2 toxin secretion (Figure 4C) and therefore display a suicidal phenotype (Figure 4D). The extended N-region is therefore especially important for immunity, and presumably less for toxin secretion.

In nature, several K2 variants exist. While the sequence we use here contains three potential initiator methionine residues at positions M1, M9, and M24, another variant^18^ contains an M9V substitution (**Figure S4B)**. While the sequence used in this study is relatively tolerant to M1 mutagenesis, the variant is not and loses immunity **(Figure S4C-E**). Speculatively, the presence of several potential initiator codons could support a more robust immunity expression and could render one variant more mutation-tolerant than another.

Combining the results from the N-terminal and C-terminal truncations of the K2 precursor, which indicate that peptides of 1-47 and 18-362 residues can both confer immunity (at 1% galactose, high-expression levels), we tested whether the overlapping region of residues 18 to 47 could independently confer immunity. This peptide, together with a set of similar peptides with additional N- or C-terminal residues, was assayed for immunity to 10 a.u. of K2 toxin in varying levels of protein expression (galactose concentrations). The peptide 18-47 did however not confer K2 resistance, and analysis of the other peptides resulted in a spectrum of immunity profiles (**Figure S4F**). It appears that the residues between 18 and 47 are most critical, but that additional residues either at the N-terminal or at the C-terminal end are required for full immunity. At high protein expression levels, the additional C-terminal residues are not critical when the extra N-terminal residues are present, and vice versa, but cannot both be omitted. However, both additional N- and C-terminal residues are required for wild-type level immunity at lower protein expression levels.

This is reminiscent of a phenomenon reported for the toxin-antitoxin system YefM-YoeB^31^. The antitoxin YefM neutralizes the YoeB toxin by forming a tight complex. Both expression of residues 1-83 and 67-92 of YefM led to suppression of YoeB toxicity, but a peptide of 67-83 did not. A slightly longer peptide of 51-83 did. Potentially, such additional facultative-essential residues could be required for peptide stability, or to stabilize a protein-protein binding interface. Whether this is indicative of a similar protein-protein binding interface for the immunity protein and either the K2 toxin or another (integral membrane) protein needs to be determined.

## Discussion

The M2 dsRNA killer virus in *S. cerevisiae* encodes both the K2 toxin as well as an immunity factor that protects the producer cell, on a single ORF. Here, through a systematic deletion scan and subsequent truncation constructs, we show that an N-terminal 49 amino acid peptide is necessary and sufficient to confer resistance to the K2 toxin in a concentration-dependent manner. Bioinformatic analyses indicate that this peptide contains a predicted transmembrane region and it is likely that this peptide also functions as the signal sequence of the K2 toxin. The immunity domain and the toxin domain (modified to contain a conventional signal peptide) are physically separable and can be expressed *in trans*. These results indicate that the K2 signal peptide plays dual roles in toxin secretion and self-protection.

Several hypotheses on the mechanism of immunity exist. For killing of a target cell, it has been proposed that K2 first binds to the fungal cell wall via β-1,6-glucans to then find a second – yet unknown – receptor in the cell membrane. The toxin subsequently disrupts membrane function. Killer toxins K1 and K2 both bind β-1,6-glucans in the cell wall, but producer cells do not show cross-resistance. Therefore, immunity is likely not mediated at the level of cell-wall β-1,6-glucan binding (i.e. via reduced levels of β-1,6-glucans or their chemical modifications). Since the peptide is present in the secretory pathway along with a potential membrane toxin receptor, it could potentially interact with (i.e. mask) such receptor before mature extracellular toxin gets a chance to bind. Alternatively, the immunity peptide could directly or indirectly interact with the external K2 toxin at the plasma membrane level. The transmembrane domains of the immunity peptides indicate that it is tethered to either the endoplasmic reticulum or the cytoplasmic membrane providing a reservoir of molecules for immunity within the secretory pathway or at the site of K2 action – the plasma membrane.

Although the signal sequences of K1, K2 and K28 all play some role in immunity, the specific amino acid sequence-dependency differs. For both K1 and K28 it has been shown regions downstream of the signal sequence are critical, more specifically the α-subunit with a C-terminal extension^32,33^. In the case of K28, a posttranslational translocation signal ensures sufficient cytoplasmic precursor levels since these precursors form complexes with and inactivate the incoming K28 toxins, moreover, this signal peptide can even be deleted without loss of immunity but cannot be replaced by signal sequences that direct the precursor into the secretory pathway in a highly efficient cotranslational fashion, which bypasses cytoplasmic precursors^32^. In contrast, the K1 precursor requires entry into the secretory pathway in order to achieve immunity^34^, but targeting does not appear to depend on the specific signal peptide sequence as studies indicate that this sequence is largely replaceable^35–37^. K2 producers are uniquely dependent on the K2 signal sequence for immunity, and the immunity and toxin domain-organization is truly modular which seems not the case for K1 and K28.

While most signal peptides do not serve additional functions and are presumably rapidly degraded, it has become evident that signal peptides can remain in the cell as stable peptides to serve post-targeting functions^38–40^. While some such signal peptides may remain in the ER membrane, other signal peptides may leave the membrane and end up in other cellular compartments^40–43^. Or, for example, signal peptides are further processed into small peptide pheromones that are involved in intercellular signaling in bacteria^44,45^. Interestingly, many of the multifunctional signal peptides that have been studied come from viruses, and are important in the viral lifecycle or contribute to immune system evasion^46–53^. RNA viruses like M2 are known for relatively compact genomes, and genome compression can be reached by use of overlapping reading frames or multifunctional proteins ^54,55^. This is especially the case for those packaged in icosahedral capsids (in contrast to those with flexible capsids), as such capsids are potentially posing a physical constraint on genome size^56–58^. The fact that killer viruses further use genome segmentation packaged into separate capsids (helper and killer virus) could also be a phenomenon evolved to relieve packaging constraints. K2 appears to be an extreme case of gene overlap, where the immunity domain takes over part of the function of the K2 toxin domain - the signal sequence – thus coupling gene expression of the toxin and the antitoxin.

The minimal identified K2 immunity factor is a small peptide with one predicted transmembrane domain. We found several reports of small proteins with similar properties in gram-positive as well as gram-negative bacteria, that protect bacteriocin producers against membrane-acting bacteriocins^59,60^. The key difference is that these immunity proteins are encoded on separate ORFs within a bacteriocin operon. The list includes immunity proteins that provide protection against a structurally divers range of bacteriocins, including lantibiotics^61^, two-peptide bacteriocins^62–64^, head-to-tail cyclized bacteriocins^65,66^, sactibiotics ^67^, microcins^68,69^ and others^70–72^. In particular, CbnZ (42 aa), which confers resistance to the two-peptide bacteriocin carnobacteriocin XY in *Carnobacterium piscicola*^62^, is a small single-transmembrane protein much like the identified minimal K2 immunity peptide.

While it is assumed that these immunity proteins localize to the membrane, where the toxins act, many questions are still open on how the toxins form membrane pores, and how small transmembrane proteins prevent or neutralize this toxicity. This represents a current gap in our understanding of how small immunity proteins confer resistance to membrane-acting protein toxins. Small proteins have historically been overlooked, but it is becoming evident that small membrane proteins have a wide range of functional potential, and can act as regulators of other integral membrane proteins and influence various cellular processes^73^. Speculatively, signal sequences can provide a source of novel functionally useful hydrophobic peptides, as signal peptides are highly abundant and tremendously sequence diverse. Signal peptide sequences could provide a reservoir of substrates for evolution, and sequence elements that do not disrupt targeting but fulfill other purposes could arise and be maintained by selection over time, including resistance to antimicrobial compounds. Alternatively, two separate ORFs could merge over time, where a hydrophobic immunity protein takes over the signal peptide function and toxin-antitoxin expression becomes linked.

Future studies should focus on elucidating the mechanism of killing of the K2 toxin, the toxin protein structure, and the mechanism the immunity peptide to protect the cell against K2 action. This includes investigation of potential interactions with the immunity protein with the K2 toxin or other proteins in the yeast membrane. While we defined a minimal immunity factor, the cleavage site and hence the size of the N-terminal immunity peptide *in vivo* is unclear. All in all, yeast killer viruses hold elegant, efficient and fascinating strategies for circumventing the lethal effects of the produced killer toxins.

## Methods

### Materials

Media and buffer components were obtained from BD Bioscience (Franklin Lakes, NJ, USA) and Sigma Aldrich (Darmstadt, Germany). Sterile, transparent round-bottom microtiter plates were obtained from Corning (Corning Inc.). Optical density measurements were performed in a SynergyMx or Powerwave (Biotek) plate reader at OD_600_.

### Strains, Media and Growth Conditions

*Escherichia coli* DH5α was used for plasmid construction and maintenance. Culturing was performed in Luria-Broth (LB), supplemented with ampicillin (100 µg/mL), kanamycin (50 µg/mL) or chloramphenicol (25 µg/mL) (37°C, 200 rpm, 16 h). *Saccharomyces cerevisiae* BY4741^74^ (*MATa leu2*Δ*0 met15*Δ*0 ura3*Δ*0 his3*Δ*1*) (derived from ATCC) was used for yeast experiments. A BY4741 Δ*PBS2* strain was obtained from the yeast deletion collection and provides an increased sensitive strain for halo assay experiments^30^. Yeast cultures were grown in YPD before transformation. After transformation, transformants were selected and cultivated on minimal SD-AS/Urea media as described elsewhere^75^ lacking histidine and/or uracil. For strain selection and maintenance 2% glucose was used as the carbon source in unbuffered minimal media. The K2 toxin is most active in acidic conditions, at pH 4.6 and 25°C. For assay conditions, cells were grown in non-inducing minimal media (2% sucrose as a carbon source, buffered with 0.5x of a citric acid/phosphate buffer at pH 4.6 as described earlier^75^), or in inducing minimal media (non-inducing media supplemented with up to 2% galactose), and cultured at 25°C, either in glass culture tubes or arrayed into 96-well plates.

| Species | Type |
| --- | --- |
| *Saccharomyces cerevisiae* | BY4741<sup>74</sup> |
| *Saccharomyces cerevisiae* | BY4741 Δ*PBS2*<sup>30</sup> |
| *Escherichia coli* | DH5α |

### Plasmid Construction

Plasmids used in this study are listed in **Supplementary Table S1** and oligonucleotides used in this study are listed in **Supplementary Table S2**. The MoClo-Yeast ToolKit^29^ was used as a cloning system to assemble constructs. Source DNA for parts was generated *via* PCR, or DNA was synthesized (gBlocks) by Integrated DNA Technologies (IDT) where necessary. For PCR amplification of new sequences, Phire Green Hot Start II PCR Master Mix (Thermo Fisher, #F126L) was used according to manufacturing instructions. Golden Gate reactions were carried out according to the standard protocol described in Lee *et al*^29^. Enzymes BsaI-HF®v2 (#R3733S) and BsmBI-v2 (#R0739S) (10.000 U/mL) and T7 ligase (#M0318S) were obtained from New England Biolabs (NEB). PCR amplicons were first cloned into the pYTK001 entry vector before assembly into a composite construct using preassembled vector backbones with GFP dropouts. Entry vectors were Sanger sequenced (Macrogen) to confirm correct sequences and the assembly digest pattern was verified by a BsmBI control restriction digest and gel electrophoresis. A high-copy pRS423-type or pRS426-type vector backbone was used for efficient protein expression.

### Preparation of concentrated K2 toxin extract

*S. cerevisiae* with plasmid pRP001 expressing K2 toxin using a constitutive ADH1 promoter was cultured in minimal media with 2% sucrose as the carbon source at pH 4.6, lacking histidine and lacking any galactose which could interfere with downstream applications. Cells were grown overnight to a final OD_600_ of 6, the cells were spun down at 5000*g* for 10 min at 4°C and the supernatant was filter sterilized. The toxin activity in the supernatant was confirmed in a halo assay and the proteins in the supernatant were concentrated 100- or 200-fold using Amicon filter units with a 10 kDa cutoff. The resulting concentrated K2 supernatant was stored at 4°C until further use. We equal the concentration of toxin present in unconcentrated supernatant to 1 arbitrary unit (a.u.), such that 100-fold concentrated extract contains 100 a.u. of toxin. We determined that a final concentration of 10 a.u. in our assays results in a robust growth-difference between immune and sensitive strains **(Figure S1A)**.

### Growth Assay in Medium With K2 Toxin

Colonies were inoculated in 200 µL of non-inducing medium in a microtiter plate, in biological triplicates. The microtiter plate was covered with a breathable membrane and incubated for 24 hours at a 30°C without agitation to reach similar growth saturation for all cultures. Subsequently, cultures were resuspended and diluted at a 1:20 ratio into a fresh plate containing inducing media supplemented with galactose (generally 1% unless concentration-dependence was assayed). Growth was measured overnight in a plate reader at 25°C (high orbital shaking), with OD_600_ measurements every 30 minutes. After 24 hours, the same procedure was repeated in a new microtiter plate containing inducing medium supplemented with 10 a.u. of K2 toxin. The resulting OD_600_ absorbance values were corrected by a blank, and corrected for the saturation of the photo-sensor as described previously^75,76^.

### Halo assay

*S. cerevisiae* ΔPBS2 was used as a sensitive indicator strain and seeded in the agar at 125 µL of OD_600_ 10 per 10 mL of plate agar. Strains to test for toxin activity were concentrated to OD_600_ 10 and 5 µL was spotted on top of the agar containing the sensitive strain. Plates were incubated at 25°C for 2-3 days until halos were visible. Plates were imaged using a FUJI imager.

### Spot assay

Strains to test for suicidal phenotypes were concentrated to OD_600_ 10 and a 1:10 dilution series in sterile water (∼10^7^ - 10^1^ cells) was spotted on top of either inducing (2% sucrose, 1% galactose) or repressing (2% glucose) agar lacking the relevant selection markers histidine and/or uracil. Plates were incubated at 25°C (inducing plates) or 30°C (non-inducing plates) for 3 days. Plates were imaged using a FUJI imager.

## Availability of data and materials sections

All data supporting the findings reported herein are enclosed in the manuscript or can be provided by the authors upon reasonable request.

