## Supporting information for "The signal sequence of yeast killer toxin K2 confers producer self-protection and allows conversion into a modular toxin-antitoxin system"

### Supplementary Figures

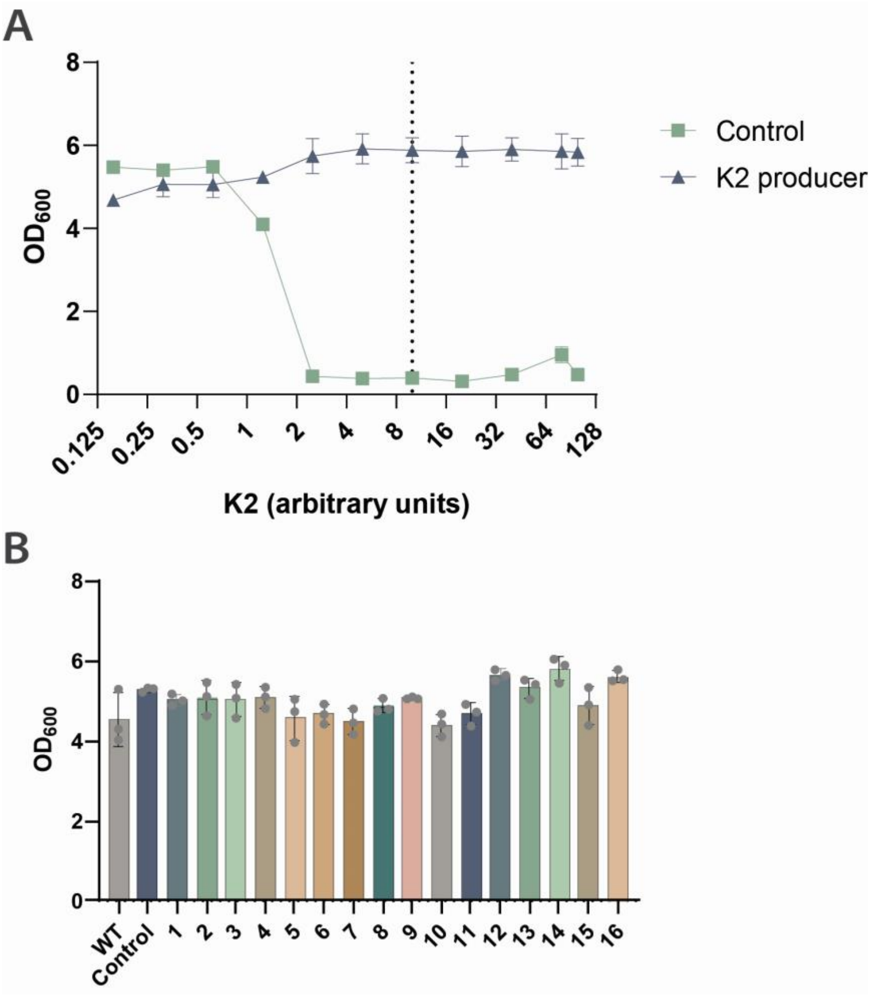


**Figure S1. Significance of the N-terminal region in K2 immunity. A)** Sensitivity of strains to varying levels of K2 toxin extract. The final $OD_{600}$ values after 24 hours of incubation in media supplemented with the respective K2 concentrations are presented. The values represent the mean of biological triplicates, with error bars indicating ± 1 standard deviation. Sensitive *S. cerevisiae* strains (Control) are inhibited above 1 a.u. of K2 toxin whereas K2 producers can be immune up to high levels of the toxin (maximum tested here is 100). The vertical dotted line marks 10 a.u., the concentration used the assays within this study. **B)** Growth of deletion constructs in inducing media without externally added K2 toxin is similar, indicating that construct expression itself did not already affect growth significantly after 24 hours. Bars represent the mean of biological triplicates, with error bars indicating ± 1 standard deviation, and individual replicate values are displayed as dots. WT: Wild-type K2, Control: Empty vector.

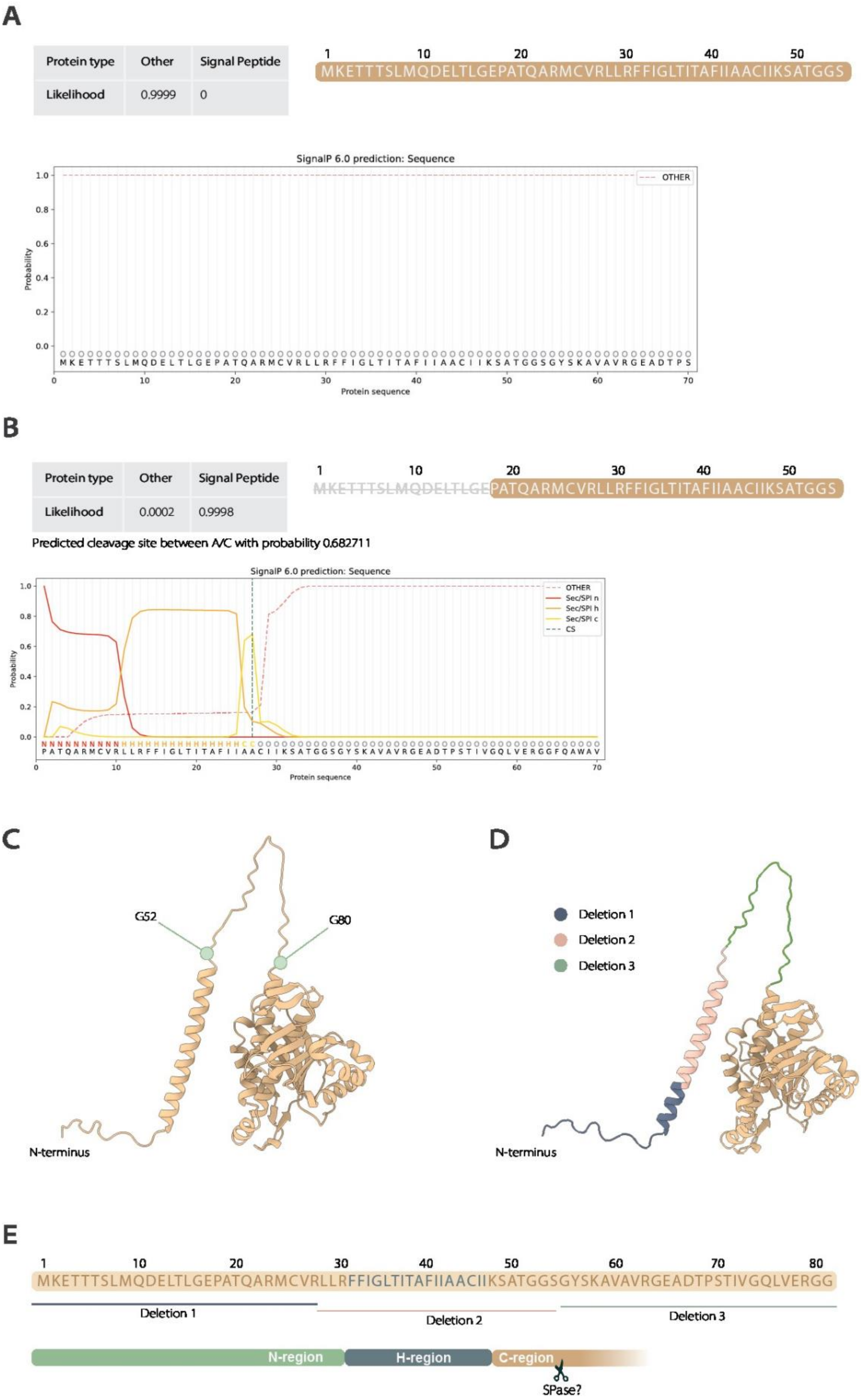
A
Protein type | Other | Signal Peptide
Likelihood | 0.9999 | 0
1 10 20 30 40 50
MKETTTSLMQDELTLGEPATQARMCVRLLRFFIGLTITAFIIAACIIKSATGGS
SignalP 6.0 prediction: Sequence
OTHER
Probability
Protein sequence
B
Protein type | Other | Signal Peptide
Likelihood | 0.0002 | 0.9998
Predicted cleavage site between A/C with probability 0.682711
1 10 20 30 40 50
MKETTTSLMQDELTLGEPATQARMCVRLLRFFIGLTITAFIIAACIIKSATGGS
SignalP 6.0 prediction: Sequence
OTHER
Sec/SPI n
Sec/SPI h
Sec/SPI c
CS
Probability
Protein sequence
C
G52
G80
N-terminus
D
Deletion 1
Deletion 2
Deletion 3
N-terminus
E
1 10 20 30 40 50 60 70 80
MKETTTSLMQDELTLGEPATQARMCVRLLRFFIGLTITAFIIAACIIKSATGGSGYSKAVAVRGEADTPSTIVGQLVERGG
Deletion 1
Deletion 2
Deletion 3
N-region
H-region
C-region
SPase?

**Figure S2**. **The vital role of the predicted K2 signal peptide in immunity. A)** When the original amino acid sequence of the K2 precursor is analyzed for a signal peptide by SignalP-6.0[1], no signal peptide is predicted. **B)** Upon the *in silico* removal of the first N-terminal 17 amino acids, a signal peptide is predicted with high confidence. Signal peptides with extended N-regions are not always recognized by bioinformatic prediction tools. Truncations of 14, 15, and 16 residues lead to a weakly predicted signal sequence (a likelihood of 0.28, 0.40 and 0.58 respectively). Earlier truncations show no prediction of a signal peptide similar to the full N-terminal sequence. **C)** The AlphaFold[2] structure prediction of the K2 precursor displays a separate N-terminal domain, consistent with the expected characteristics of signal peptides that are not part of the mature secreted protein. The glycine residues at positions 52 and 80 flank a loop or linker region situated between what is predicted to be the alpha-helical H-region of the signal peptide and the downstream toxin domain, which likely contains a protease processing site for releasing the toxin domain. **D)** The regions that were deleted in the loss-of-resistance constructs are indicated (see also Figure 1 in the main text). While deletion 1 and 2 delete part of the N-region and H-region of the predicted signal peptide, the region targeted by deletion 3 forms the loop region connecting the H-region of the signal peptide and the presumed toxin domain. Speculatively, this loop may play a significant role in proper folding and/or processing of the precursor into a functional immunity and toxin domain. **E)** Overview of the predicted N-region, H-region and C-region, along with the deletion segments 1, 2 and 3 within the N-terminal amino acid sequence of the K2 precursor.

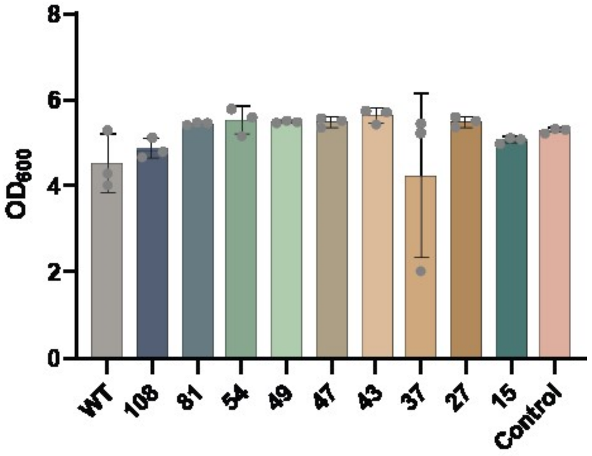


**Figure S3. Growth in inducing media in absence of externally added toxin.** Growth of truncation constructs in inducing media without externally added K2 toxin is similar, indicating that construct expression itself did not already affect growth significantly after 24 hours. Bars represent the mean of biological triplicates, with error bars indicating ± 1 standard deviation, and individual replicate values are displayed as dots. WT: Wild-type K2, Control: Empty vector.

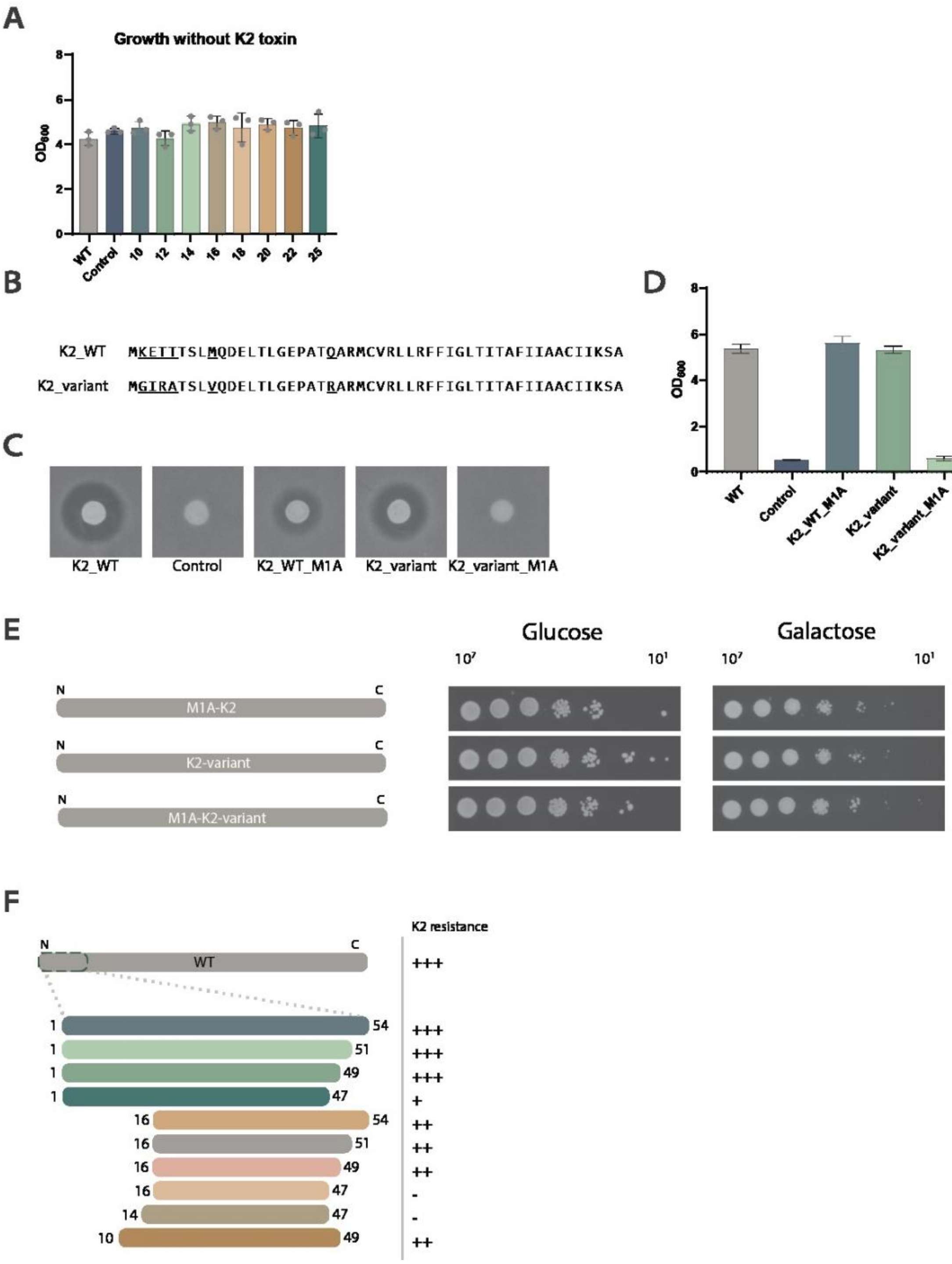


**Figure S4. Characteristics of the N-terminal peptide. A)** Growth of constructs in inducing media without added K2 toxin. Final $OD_{600}$ values after 24 hours are displayed. Bars represent the mean of three biological replicates with error bars representing +- 1 standard deviation. Individual replicates are depicted as dots. **B)** The N-terminal sequences of the K2 variant used in this study (K2_WT) and a K2 variant published elsewhere[3] (K2_variant) show several amino acid substitutions (underscored). Specifically, K2_WT contains three potential methionine initiator codons (M1, M9, M24), whereas K2_variant lacks M9. **C)** Halo assay on a sensitive background strain to determine whether active toxin is secreted. The first methionine residue was mutated into alanine, which results in lack of toxin expression of K2_variant, whereas K2_WT still retains partial toxin secretion. **D)** While both wild-type sequences confer immunity to K2, only K2_WT retains immunity after the M1A mutation. The K2_variant shows a loss-of-resistance phenotype. **E)** Consistent with these results, none of the strains display a suicidal phenotype – K2_WT because it supports immunity and K2_variant because of simultaneous lack of toxicity and immunity. **F)** We conducted growth assays over 24 hours, with a range of different inducing galactose concentrations, for several peptide variants in presence of a concentration of 10 a.u. of toxin. Based on the resulting dynamics, we define different immunity profiles, wild-type level (+++), intermediate (++), low (+), and no immunity (-). The first three (+++, ++, +) all confer K2 resistance similarly at high-expression levels (1% galactose and above) but show different profiles at lower expression levels.

### Supplementary Notes

#### Note 1. The signal peptidase cleavage site.

After *in silico* adjustments to the N-region, different prediction tools suggest cleavage sites between residues 43 and 44, 44 and 45, or 52 and 53[1,4,5]. A cleavage site between residues 43 and 44 has also been suggested in previous studies[6]. However, based on our findings that a resulting 43 amino acid peptide would not confer immunity to the K2 toxin, and the fact that immunity is conferred in a concentration-dependent manner and thus a number of functional immunity peptides would need to be present at any given time to protect the cell, it seems an unlikely case that such cleavage, resulting in dysfunctional immunity peptides, could provide the high-level protection observed in K2 producers. Therefore, we believe that it is more likely that a cleavage site around position 52-56 (multiple potential cleavage sites present) or downstream is used *in vivo*. The Lassa virus glycoprotein signal peptide is an example of a peptide that is longer than usual (58 residues) and was predicted to be cleaved at position 34, but was experimentally shown to be cleaved only after position 58[7]. Experimental data is needed to determine how the K2 precursor is processed *in vivo*.

**Supplementary Tables**

**Supplementary Table S1.** Plasmids used in this study**.**

| Plasmid | Name | Notes |
|---|---|---|
| **General plasmids** | | |
| pRP001 | pADH1_K2_tADH1 | pRS423-type |
| pRP002 | pGAL1_K2_tADH1 | pRS423-type |
| pRP223 | pGAL1_K2_tENO1 | YTK assembly, pRS423-type |
| pRP246 | Spacer histidine marker (‘empty plasmid’) | YTK assembly, pRS423-type |
| pRP234 | Spacer uracil marker (‘empty plasmid’) | YTK assembly, pRS426-type |
| **SbfI-restriction site constructs (Figure 1)** | | |
| pRP035 | 1 | SbfI site replaces aa 1 to 27 |
| pRP036 | 2 | SbfI site replaces aa 28 to 54 |
| pRP037 | 3 | SbfI site replaces aa 55 to 81 |
| pRP038 | 4 | SbfI site replaces aa 82 to 108 |
| pRP039 | 5 | SbfI site replaces aa 109 to 135 |
| pRP040 | 6 | SbfI site replaces aa 136 to 162 |
| pRP041 | 7 | SbfI site replaces aa 163 to 189 |
| pRP042 | 8 | SbfI site replaces aa 190 to 216 |
| pRP043 | 9 | SbfI site replaces aa 217 to 243 |
| pRP044 | 10 | SbfI site replaces aa 244 to 268 |
| pRP045 | 11 | SbfI site replaces aa 269 to 284 |
| pRP046 | 12 | SbfI site replaces aa 285 to 300 |
| pRP047 | 13 | SbfI site replaces aa 301 to 316 |
| pRP048 | 14 | SbfI site replaces aa 317 to 332 |
| pRP049 | 15 | SbfI site replaces aa 333 to 348 |
| pRP050 | 16 | SbfI site replaces aa 349 to 362 |
| **Systematic C-terminal truncation constructs (Figure 3)** | | |
| pRP075 | 54 | pGAL1_aa1to54_tADH1 |
| pRP076 | 81 | pGAL1_aa1to81_tADH1 |
| pRP077 | 108 | pGAL1_aa1to108_tADH1 |
| pRP082 | 47 | pGAL1_aa1to47_tADH1 |
| pRP083 | 27 | pGAL1_aa1to27_tADH1 |

| | | |
|---|---|---|
| pRP084 | 37 | pGAL1_aa1to37_tADH1 |
| pRP085 | 15 | pGAL1_aa1to15_tADH1 |
| pRP086 | 43 | pGAL1_aa1to43_tADH1 |
| pRP162 | 49 | pGAL1_aa1to49_tADH1 |
| **Systematic N-terminal truncation constructs (Figure 4)** | | |
| pRP228 | 10 | pGAL1_aa10to362_tENO1 |
| pRP247 | 12 | pGAL1_aa12to362_tENO1 |
| pRP248 | 14 | pGAL1_aa14to362_tENO1 |
| pRP249 | 16 | pGAL1_aa16to362_tENO1 |
| pRP250 | 18 | pGAL1_aa18to362_tENO1 |
| pRP251 | 20 | pGAL1_aa20to362_tENO1 |
| pRP252 | 22 | pGAL1_aa22to362_tENO1 |
| pRP232 | 25 | pGAL1_aa24to362_tENO1 |
| **K2 resistance peptide constructs (Figure S4)** | | |
| pRP241 | 1 to 54 | pGAL1_aa1to54_tENO1 |
| pRP230 | 1 to 47 | pGAL1_aa1to47_tENO1 |
| pRP245 | 1 to 51 | pGAL1_aa1to51_tENO1 |
| pRP243 | 1 to 49 | pGAL1_aa1to49_tENO1 |
| pRP222 | 10 to 47 | pGAL1_aa10to47_tENO1 |
| pRP273 | 10 to 49 | pGAL1_aa10to49_tENO1 |
| pRP270 | 14 to 47 | pGAL1_aa14to47_tENO1 |
| pRP271 | 16 to 47 | pGAL1_aa16to47_tENO1 |
| pRP272 | 16 to 54 | pGAL1_aa16to54_tENO1 |
| **Constructs tested for suicide phenotype (Figure 2 and 3, Figure S4)** | | |
| pRP233 | 1 to 47 | Alike pRP230 but with a uracil marker for co-transformation |
| pRP187 | pPTK007-K2 | αMFΔ-K2 |
| pRP188 | pPTK014-K2 | Killer-αMFΔ-K2 |
| pRP163 | LeucineTM | leuTM-K2 |
| pRP213 | T38KFP suicide construct[17] | T38KFP-K2 |
| pRP087 | K2 M1A | Mutated initiator methionine |
| pRP088 | Gulbiniene K2 variant[18] | Original variant of K2 |
| pRP089 | Gulbiniene K2 M1A | Mutated initiator methionine |

| Yeast ToolKit (YTK) parts that were used for vector assemblies | | |
|---|---|---|
| pYTK001 | Entry vector | |
| pYTK002 | ConLS | Type 1 |
| pYTK030 | pGAL1 | Type 2 |
| pYTK048 | Type 234 spacer | Type 2-3-4 |
| pYTK051 | tENO1 | Type 4 |
| pYTK072 | ConRE | Type 5 |
| pYTK074 | URA3 | Type 6 |
| pYTK076 | HIS3 | Type 6 |
| pYTK082 | 2μ | Type 7 |
| pYTK083 | AmpR-ColE1 | Type 8 |
| pPTK007 | αMFΔ | Type 3a |
| pPTK014 | Killer-αMFΔ | Type 3a |
| pRP209 | pRS426-type backbone | Assembly vector |
| pRP278 | pRS423-type backbone | Assembly vector |

**Supplementary Table S2.** Primers used in this study.

| **Primer** | **Note** | **Sequence** |
|---|---|---|
| C-terminal truncation constructs | | |
| RP66 | K2 fw | TTAACTAATACTTTCAACATTTTCGGTTTG |
| RP91 | Position 15 rv | GTACATACATAAACATACGCGCACAAAAGCAGAGACAGATATCT TA CAGGGTCAGTTCATCCTGC |
| RP89 | Position 27 rv | GTACATACATAAACATACGCGCACAAAAGCAGAGACAGATATCT TA GCGCACGCACATGC |
| RP90 | Position 37 rv | GTACATACATAAACATACGCGCACAAAAGCAGAGACAGATATCT TA AATGGTCAGGCCAATAAAAAAG |
| RP96 | Position 43 rv | GTACATACATAAACATACGCGCACAAAAGCAGAGACAGATATCT TA CGCAATAATAAACGCGGTAATG |
| RP88 | Position 47 rv | GTACATACATAAACATACGCGCACAAAAGCAGAGACAGATATCT TA AATAATGCACGCCGCAATAATAAAC |
| RP116 | Position 49 rv | GTACATACATAAACATACGCGCACAAAAGCAGAGACAGATATCT TA GCTTTTAATAATGCACGC |
| RP62 | Position 54 rv | GTACATACATAAACATACGCGCACAAAAGCAGAGACAGATATCT TA GCTGCCGCCGGTCG |
| RP63 | Position 81 rv | GTACATACATAAACATACGCGCACAAAAGCAGAGACAGATATCT TA GCCGCCGCGTTCCAC |
| RP64 | Position 108 rv | GTACATACATAAACATACGCGCACAAAAGCAGAGACAGATATCT TA CACCGCCGCGGTCAC |

| K2 variants and suicidal constructs | | |
|---|---|---|
| RP103 | K2_WT-M1A | CCGAATTCCCAAAAGAA GCT AAA GAA ACC ACC ACC AGC CTG ATG CAG GAT GAA CTG ACC CTG GGC GAA CCG GCG ACC CAG GCG CGC ATG TGC GTG CGC CTGCTGCGCTTTTTTATTG |
| RP104 | K2_variant | CCGAATTCCCAAAAGAA ATGGGTATTAGAGCTACCAGCCTGGTTCAGGATGAACTGACCCT GGGCGAACCGGCGACCAGAGCGCGCATGTGCGTGCGC CTGCTGCGCTTTTTTATTG |
| RP105 | K2_variant-M1A | CCGAATTCCCAAAAGAA GCTGGTATTAGAGCTACCAGCCTGGTTCAGGATGAACTGACCCT GGGCGAACCGGCGACCAGAGCGCGCATGTGCGTGCGC CTGCTGCGCTTTTTTATTG |
| RP117 | LeuTM construct | GCATGTGCGTGCGC CTGCTGTTGTTGTTGTTGTTGCTGTTGTTGTTGTTGTTGTTGTTGT TGTTGTTGTTGTTGAAAAGCGCGACCGGCGGCAGC GGCTATAGCAAAGCGGT |
| gBlock7 | T38KFP construct | GAATTCCCAAAAGAAATGAAAGAAACCACCACCAGCCTGATGC AGGATGAACTGACCCTGGGCGAACCGGCGACCCAGGCGCGCAT GTGCGTGCGCCTGCTGCGCTTTTTTATTGGCCTGACCATTAAATT TCCGGCGTTTATTATTGCGGCGTGCATTATTAAAAGCGCGACCGG CGGCAGCGGCTATAGCAAAGCGGTGGCGGTGCGCGGCGAAGCG GATACCCCGAGCACCATTGTGGGCCAGCTGGTGGAACGCGGCGG CTTTCAGGCG |
| SB73 | gBlock 7 fw | GCACAATATTTCAAGCTATACCAAGCATACAATCAACTGAATTC CCAAAAGAAATGAAAG |
| SB74 | gBlock 7 rv | TTTTCGCAAACAGATAAATGCCCGCGCCCACCGCCCACGCCTGA AAGCCGCCGC |
| N-terminal truncation constructs, signal peptide substitution and peptide variants | | |
| SB160 | K2 fw (type 3) | GCATCGTCTCATCGGTCTCATATGAAAGAAACCACCAC |
| SB161 | K2 rv (type 3) | ATGCCGTCTCAGGTCTCAGGATTTAGCCGCTATCGC |
| RP126 | K2 54-362 fw (type 3b) | GCATCGTCTCATCGGTCTCATTCTAGCGGCTATAGCAAAG |
| RP258 | Position 10 fw (type 3) | GCATCGTCTCATCGGTCTCA T ATGCAGGATGAACTGACC |
| RP263 | Position 12 fw (type 3) | GCATCGTCTCATCGGTCTCATATGGAACTGACCCTGG |
| RP264 | Position 14 fw (type 3) | GCATCGTCTCATCGGTCTCATATGACCCTGGGCGAAC |
| RP265 | Position 16 fw (type 3) | GCATCGTCTCATCGGTCTCATATGGGCGAACCGGCG |
| RP266 | Position 18 fw (type 3) | GCATCGTCTCATCGGTCTCATATGCCGGCGACCCAG |
| RP267 | Position 20 fw (type 3) | GCATCGTCTCATCGGTCTCATATGACCCAGGCGCG |
| RP288 | Position 22 fw (type 3) | GCATCGTCTCATCGGTCTCATATGGCGCGCATGTGC |
| RP262 | Position 25 fw (type 3) | GCATCGTCTCATCGGTCTCAT ATGTGCGTGCGC |
| RP259 | Position 47 rv (type 3) | ATGCCGTCTCAGGTCTCAGGAT TTA AATAATGCACGCCGCAAT |
| RP271 | Position 49 rv (type 3) | ATGCCGTCTCAGGTCTCA GGAT TTA GCTTTTAATAATGCACGC |

| | | |
|---|---|---|
| RP272 | Position 51 rv (type 3) | ATGCCGTCTCAGGTCTCA GGAT TTA GGTCGCGCTTTTAATAATG |
| RP270 | Position 54 rv (type 3) | ATGCCGTCTCAGGTCTCA GGAT TTA GCTGCCGCCGG |
| Sequencing primers | | |
| RP46 | Sequencing pYTK001 fw | TTACGGTTCCTGGC |
| RP47 | Sequencing pYTK001 rv | TGATAGATCCAGTAATGACC |
| RP01 | Sequencing pRP002 fw | CTT TCA ACA TTT TCG GTT TG |
| RP02 | Sequencing pRP002 rv | GCTTAAACACGTCTTTTCC |
| SB181 | Sequencing assemblies fw | CACAGACATTAACCCACAG |
| SB159 | Sequencing assemblies rv | GTCAGTGTGAGCACCAC |
